# Tropomyosin in mugwort cross-reacts to house dust mite, eliciting non-Th2 response in allergic rhinitis patients sensitized to house dust mite

**DOI:** 10.1101/2020.10.13.338533

**Authors:** Feng Lan, Limin Zhao, Su Duan, Nan Zhang, Haibo Zhang, Ming Zheng, Qiqi Wang, Xu Zhang, Xiangdong Wang, Sun Ying, Claus Bachert, Luo Zhang

**Author notes:** **Corresponding authors:** Prof. Luo Zhang, Beijing Institute of Otolaryngology, Department of Otolaryngology-Head and Neck Surgery, Capital Medical University, No. 17, HouGouHuTong, Dongcheng District, Beijing 100730, China. These authors contributed equally to the study (Feng Lan, Limin Zhao).

## Abstract

**Background:** Mugwort and house dust mite (HDM) are two of the most common inhalant allergens in Asia; however, whether or not mugwort affects polysensitized HDM^+^ allergic rhinitis (AR) patients has not been elucidated.

**Methods:** Overall, 15884 AR outpatients were assessed for clinical status. Amino acid sequences of mugwort were determined by mass spectrometry. Afterward, cross-reactivity between mugwort tropomyosin and Dermatophagoides pteronyssinus 10 (Der p10) was analysed by ELISA inhibition and basophils activation experiments. To compare immunologic responses eliciting by two different tropomyosins, peripheral blood mononuclear cells (PBMCs) of HDM-monosensitized patients were stimulated by mugwort, HDM, Der p10 and synthetic peptides representing mugwort tropomyosin respectively.

**Results:** Polysensitized HDM^+^AR patients were mainly sensitized to cat and mugwort, and the positive rate of monosensitized HDM^+^AR out-clinic patients was increased during the mugwort pollen season. Mugwort tropomyosin protein had similar structural domains to HDM tropomyosin, Der p10. ELISA inhibition experiment showed synthetic mugwort tropomyosin peptide inhibited IgE binding to Der p10; mugwort tropomyosin peptide activated basophils which were primed by HDM-specific IgE. Unlike HDM and Derp 10, mugwort and mugwort tropomyosin mainly induced IFN-γ and IL-17, release in PBMCs of monosensitized HDM^+^AR patients, but not IL-5.

**Conclusions:** Pan-allergen tropomyosin is a major protein accounting for the cross-reactivity between mugwort and HDM, which reminds HDM^+^ patients to reduce mugwort exposure in mugwort pollen season in virtue of the tropomyosin induced mild inflammation.

## INTRODUCTION

Allergic rhinitis (AR) is an upper airway allergic inflammatory disease, which causes symptoms of sneeze, runny nose, nasal obstruction and itchy nose, and is mediated by type 2 helper (Th2) cells and immunoglobulin E (IgE)^1,2^. Among the common triggering allergens, house dust mites (HDM), mould spores and animal dander mainly cause symptoms of perennial allergic rhinitis, whereas, a large variety of pollens from different geographical regions contributes to symptoms of seasonal allergic rhinitis^3^. Some AR patients are found to be polysensitized to more than one allergen^4^, and an increasing number of sensitizations strongly predisposes AR patients to allergic asthma^5,6^. Thus, the treatment for polysensitized AR patients has a close relationship with asthma management^7^.

Allergen specific immunotherapy (AIT) is an effective therapeutic method for monosensitized AR patients^7^. However, management approaches to polysensitized AR patients by AIT are not standardized yet. There are intercontinental differences in allergen products available for AIT in polysensitized patients^8^. Desensitization to the most clinically relevant allergen is often used to treat polysensitized patients in Europe and in China, while mixtures of extracts are recommended in the United States^9,10^. Differences in therapeutic effects of single allergen-specific immunotherapy have been shown: more effective in reducing the symptoms are observed in those of monosensitized patients than that of polysensitized patients treated with the same dose^11,12^; no obvious change in HDM-specific IgE production and a lower concentration of HDM-specific IgG4 is noticed in polysensitized patients compared with those of monosensitized patients after AIT^12,13^. Polysensitization is mainly caused by cross-reactivity among closely related allergens, or allergens from other sources. The identification of primary causal allergen(s) and sensitization to cross-reacting allergens helps us find efficient ways to treat polysensitized AR patients in the near future.

HDM and mugwort have been regarded as the two most common and clinically relevant sensitizing allergens in AR patients in Asia ^14^. HDM cross-reacts with allergens from other invertebrates, including other species of mites, insects, mollusks, and crustaceans^15^. It is not clear whether or not there is cross-reactivity between HDM and mugwort; consequently, whether or not mugwort affects polysensitized HDM^+^AR patients. Thus, it would clearly be of interest to explore whether HDM and mugwort share similar structural features that elicit a common immunologic response in polysensitized and monosensitized patients. In view of this, the present study has specifically investigated cross-reactivity between HDM and mugwort in HDM^+^AR patients.

## MATERIAL AND METHODS

### Study design and subjects

Subjects with AR based on criteria of the Allergic Rhinitis and its Impact on Asthma (ARIA) consensus statement^16^ were recruited consecutively from the allergy-rhinology outpatient clinic of Beijing Tongren Hospital. On recruitment, each subject completed a questionnaire to record demographic data, nasal symptom severity, and history of asthma; and blood samples were collected from each subject for analysis of serum specific IgE antibodies. Peripheral mononuclear cells (PBMCs) were also prepared from blood samples of some HDM^+^AR patients and healthy controls for this study. None of the subjects had received any allergen-specific immunotherapy or monoclonal antibody treatment. The study was approved by the Medical Ethics Committee of Beijing Tongren Hospital, and all patients provided written informed consent before entry into the study and collection of any samples.

### Serum antigen-specific IgE measurements

The presence of IgE antibodies in blood was determined using a EUROLINE Atopy Screen (DP 3713 E; Lubeck Germany), which comprised two sets of allergens; one with a mix of aeroallergens [including tree mix (willow, poplar, elm), common ragweed, mugwort, house dust mite mix (Dermatophagoides pteronyssinus (Der p), Dermatophagoides farina (Der f), house dust, cat, dog, cockroach German, mould mix (Penicillium notatum, Cladosporium herbarum, Aspergillus fumigatus, Alternaria alternata) and hops], and one with a mix of food allergens [including egg white, cow’s milk, peanut, soybean, beef, mutton, sea fish mix (codfish, lobster, scallop), shrimp, and crab]. Furthermore, concentrations of Der f2 specific IgE, Der p1 specific IgE, and total IgE were also measured using the ImmunoCAP system (Immunodiagnostics; Thermo Fisher Scientific, Uppsala, Sweden). Allergen-specific IgE >0.35 kU/L was considered as positive.

### Mass Spectrometry

Prior to analysis, 100 mg samples of mugwort (*Artemisia* annua (A. annua) and *Artemisia* sieversian (A. sieversian)) were separately prepared as peptide solutions by denaturing and treatment with protease trypsin according to the method described by León and colleagues^17^; and then analysed in a Triple-TOF 6600 mass spectrometer (Sciex, United States) fitted with a Nanospray III source (Sciex). The ion spray voltage was 2300 V, declustering potential 80 V, curtain gas 35 psi, nebulizer gas 5psi, and interface heater temperature 150 °C. The peptides were introduced into the mass spectrometer via Nona 415 liquid chromatography column (Sciex) eluted with water/acetonitrile/formic acid (buffer B: 2/98/0.1%). In this regard, samples (4 μL) were injected onto a C18 desalted column (3 μm, 120Å, 350 μm×0.5 mm), and separated onto a C18 analysis column (3 μm, 120Å, 75 μm×150 mm) with gradients ranging from 5 to 16% buffer B in the first 25 min, from 16 to 26% buffer B in the next 20 min, from 26 to 40% buffer B in the following 3 min, from 40 to 80% buffer B in the next 5 min, and finally from 80 to 5% buffer B in the final 7 mins; at a flow rate of 0.6 μL/min. The peptides present in the samples, were matched to the UniProt *Artemisia* annua databases, and identified for the corresponding proteins. A standardised preparation of common repeat peptide sequences of tropomyosin protein from both A. annua and A. sieversian (SynPeptide, Shanghai, China) was selected for further functional analysis as follows: VGSPDESYEDFTNSLPSNECR.

### Comparative modelling of three-dimensional (3D) protein structures

Homology modelling for tropomyosin of mugwort (A. annua and A. sieversian) was established on the SWISS-MODEL server. The modelling template was selected from RCSB PDB Data Bank (PDB ID: 1f7s and 1xwv) and molecular dynamics simulations were carried out for mugwort using AMBER ff14SB force field. The structures were explicitly solvated with a water box from the outermost atoms of the molecules. Periodic boundary conditions were used and the net negative charge neutralized with Na^+^ counterion. The best model was selected for the analysis of the similarity between tropomyosin of mugwort and Der p10. The structure of Der p10 was established on the SWISS-MODEL server.

### HDM-specific IgE blockage by synthesised mugwort tropomyosin peptides

Serum samples of 15 HDM^+^AR patients each with a high or low level of HDM-specific IgE were used to assess the IgE blocking potential of synthetic peptides of mugwort tropomyosin. Briefly, 200 μL of serum from HDM^+^AR patients were incubated with or without mugwort tropomyosin peptides (1000 ng/mL for each) for 1 h at room temperature, and at the end of incubation the serum samples were analysed for the concentrations of Der p1-specific IgE and Der f2-specific IgE using the ImmunoCAP system.

### Basophil activation test

PBMCs isolated from non-allergic donors, 5X10^5 cells were stripped in 2ml ice cold lactic acid buffer (0.13M KCl, 0.05M NaCl, 0.01M lactic acid, pH=3.9) for 30s as described before^18^. After washing 3 times by PBS, cells were pre-incubated with sera from HDM-allergic individuals for 1 h at 37°C. And then, cells were stimulated by different concentrations of mugwort tropomyosin or Derp 10 (50, 500 ng/mL) in hepes buffer containing IL-3 (R&D, Minneapolis, Minnesota, USA). In the meanwhile, cells exposed to FLMP (Sigma, St. Louis, USA)) were taken as a positive control. The reaction was stopped by EDTA buffer (20 mM). In the end, PBMCs were stained with basophil surface markers: CD123BV650, CCR3-APC-fire750 and CD63-PE (Biolegend, San Diego, CA, USA), and the percentages of CD63^+^CD123 ^+^CCR3^+^ cells were analysed by Flowjo software.

### ELISA inhibition experiment

Plates were pre-coated with Der p10 obtained from CUSABIO company (Wuhan, China) overnight at 4°C. Non-specific binding in the well was reduced by incubation with PBS supplemented with 1% BSA and 0.05% tween 20 for 6 h at room temperature. Inhibition was performed by adding sera from HDM monosensitzed patients with synthetic mugwort tropomyosin peptides (50, 500 ng/mL), and sera without peptides were taken as non-inhibition conditions. Anti-human IgE (2 ug/mL, NOVUS, USA) were added, followed by streptavidin-HRP conjugated secondary antibodies (diluted 1:2,000; EasyBio, Beijing, China). Absorbance at 405 nm was determined using an ELISA reader (BioTek, Vermont, USA). All experiments were performed in duplicate. Percent inhibition was calculated using the following equation: percent inhibition = 100- [(OD of serum with tropomyosin peptide /OD of serum without peptide) × 100].

### Stimulation of PBMCs *ex vivo*

PBMCs were isolated from the blood of 6 healthy and 12 AR donors using Ficoll-Hypaque density gradient centrifugation according to the standard protocol (Lymphoprep^TM^, Nycomed Pharma, Oslo, Norway). Cells were plated at a density of 1×10^6^ cells/well in a 24-well plate in 0.5 mL RPMI 1640 (Gibco, USA) culture medium alone, or with HDM (Der p1 extract; 0.2, 1, 5 μg/mL; GREER Laboratories, Lenoir, NC, USA), mugwort (1, 10, 100, 1000 ng/mL; locally prepared in Beijing Tongren Hospital), or mugwort tropomyosin peptides in the culture medium, and then incubated at 37 °C in 5% CO_2_ for 48 h. At the end of incubation, the cell suspensions were collected and the supernatants were assessed for IL-5, IL-17, and IFN-γ using Luminex xMAP suspension array technology in a Bio-Plex 200 system (Bio-Rad, MI). All cytokine kits were purchased from R&D Company and the cytokines were expressed as pg/mL.

### Statistical analysis

Statistical analysis was performed using the SPSS version 22.0 software package (IBMCorp, Armonk, NY, USA). Categorical variables were described using frequencies and/or percentages and continuous variables were presented as means ± standard deviation (SD). Multiple logistic regression was used to analyse the possible risk factors for polysensitized HDM^+^AR patients. The influence of polysensitization on asthma development was assessed by the Chi-squared test. The prevalence of different allergens in HDM^+^AR patients was estimated using Fisher’s exact test and logistic regression. The Wilcoxon test was used for paired between-group comparisons of the effect of specific antigen stimulation on the release of cytokines from PBMCs, and the effect of mugwort peptides on blocking Der p1 and Der f2 sIgEs. *P* values of less than 0.05 were regarded as statistically significant.

## RESULTS

### Mugwort may affect the prevalence of HDM+AR patients

A total of 497 HDM^+^AR patients were recruited into the study. Overall, 64.6% of the HDM^+^AR patients were monosensitized, and 35.4% polysensitized (**Table 1**). Comorbid asthma was more prevalent in 19.32% of all polysensitized AR patients compared to in 9.03% of all monosensitized AR patients (*p* = 0.001). Type of sensitizing allergens was further analysed in polysensitized HDM^+^AR patients in parallel with the HDM-specific IgE level. Regardless of the level of HDM-specific IgE detected, sensitization was greatest to inhalant allergens in the polysensitized HDM^+^AR patients (**Figure 1A**). The five most prevalent inhalant allergens in the polysensitized HDM^+^AR patients were cat (27.8%, 95% CI:21.2%-34.5%), mugwort (26.1%, 95% CI:19.6%-32.7%), house dust (21.6%, 95% CI:15.5%-27.7%), cockroach (20.5%, 95% CI:14.4%-26.5%) and hops (10.8%, 95% CI: 6.2%-15.4%) (**Figure 1B**). The number of monosensitized HDM^+^AR patients was increased from July to August, and the number of polysensitized HDM^+^AR patients from July to September (**Figure 1C**); which appeared to follow the trend of the mugwort pollen season seen from July to early September in 2018 ^19^.

**Table 1.** Demographic and clinical characteristics of house dust mite-positive allergic rhinitis patients investigated.

|  | Monosensitization<br>(n=321) (64.4%) | Polysensitization<br>(n=176) (35.4%) | P value | Odds<br>ratio | 95%CI |
| --- | --- | --- | --- | --- | --- |
| **Gender** |  |  | 0.058 | 0.690 | 0.470-1.012 |
| Male | 158 (49.22) | 101 (57.39) |  |  |  |
| Female | 163 (50.78) | 75 (42.61) |  |  |  |
| Age | 29.60±11.55 | 28.77±12.09 | 0.449 |  |  |
| Family history of AR | 69 (21.50) | 54 (30.68) | 0.056 | 1.522 | 0.989-2.342 |
| Smoking and drink | 222 (69.16) | 99 (56.25) | 0.003* | 0.560 | 0.381-0.823 |
| Co-morbid Allergic status |  |  |  |  |  |
| Asthma | 29 (9.03) | 34 (19.32) | 0.001* | 2.486 | 1.448-4.267 |
| Atopic Dermatitis | 28 (8.72) | 24 (13.64) | 0.118 | 1.506 | 0.818-2.772 |
| Allergic conjunctivitis | 34 (10.59) | 17 (9.66) | 0.206 | 0.648 | 0.331-1.269 |
| **HDM specific IgE (kU/L)** | 25.44±26.66 | 29.89±29.92 | 0.112 | 1.005 | 0.999-1.012 |

\*P< 0.05, HDM= house dust mite.

**Figure 1.**
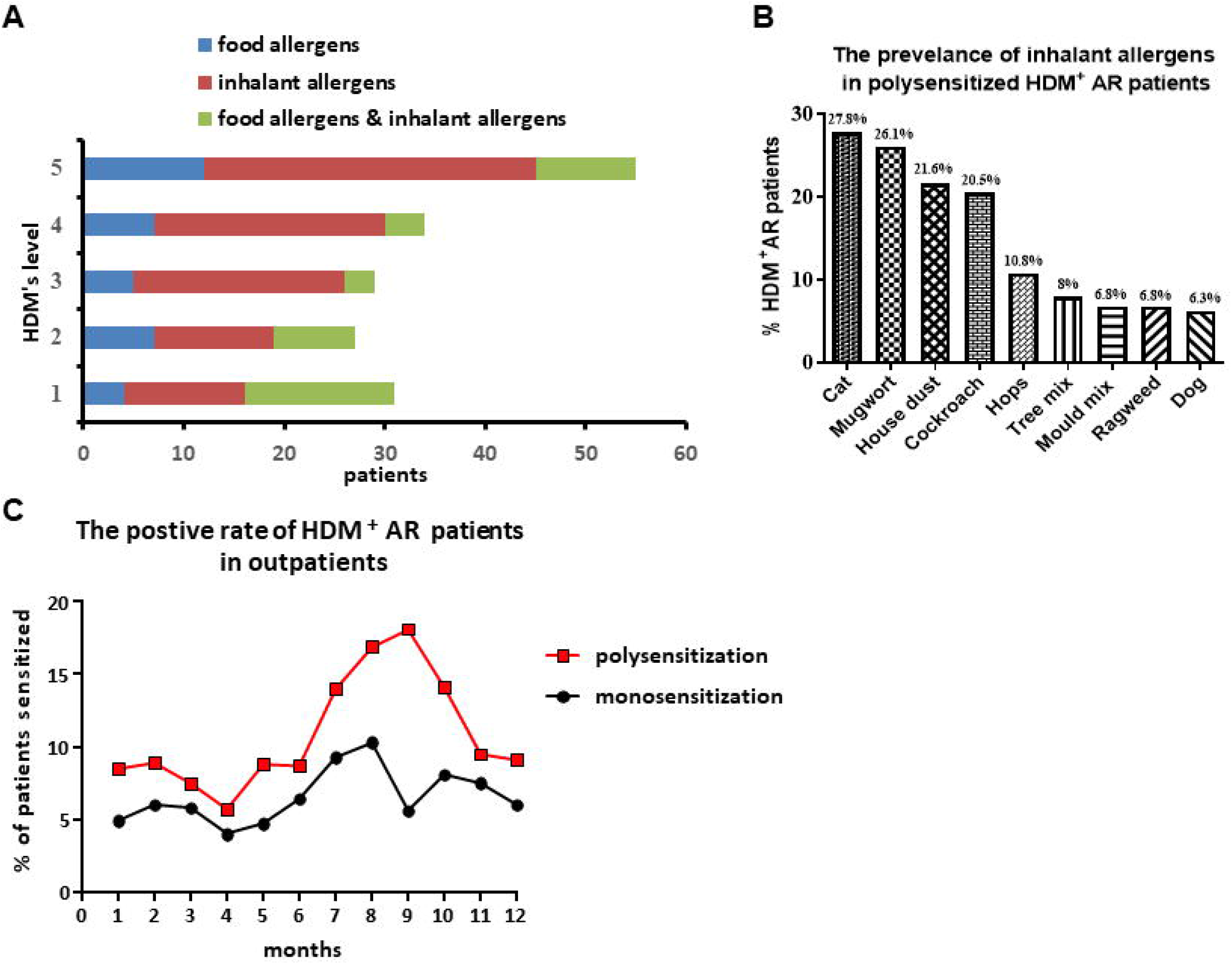
**A**: The distribution of allergen types, according to the concentrations of house dust mite (HDM) specific IgE in polysensitized HDM^+^ allergic rhinitis (AR) patients. Food allergens included crab, shrimp, soybean, sea fish mix 1, egg white, beef, cow’s milk, peanut and mutton; while inhalant allergens included cat, mould mix 1, mugwort, hops, common ragweed, dog, cockroach, German, tree mix 2 and house dust. **B**: The prevalence of inhalant allergens in polysensitized HDM^+^AR patients (n=176). **C**: The positive rate of monosensitized and polysensitized HDM^+^AR patients in recruited outpatients from January 2018 to December 2018 (n=15354).

### Tropomyosin was involved in mugwort-HDM cross-reactivity due to the similar structure

Alignment of peptide amino acid sequences of A. annua and A. sieversian allergens in UniProt *Artemisia* annua databases indicated the presence of cross-reactivity protein tropomyosin (**Table 2**). Similarly, the presence of cross-reactivity proteins profilin and lipid transfer protein were also found in both A. annua and A. sieversian (data not shown). However, alignment analysis of the amino acid sequences demonstrated for the tropomyosin proteins in mugwort with the amino acid sequences for Der p10 (a member of the tropomyosin group protein in HDM), showed that there was no homology. Comparative 3D-modeling of the tropomyosin protein structures of mugwort *vs* Der p10, optimized using AMBER14 molecular dynamics software, demonstrated that the tropomyosin proteins in mugwort contained α-helices and β-sheets, which were comparable to Der p10 (**Figure 2A**).

**Table 2:** Amino acid sequences of Artemisia annua and Artemisia sieversian tropomyosin fragments detected by mass spectrometry.

| Mugwort |  |  |  |
| --- | --- | --- | --- |
| Species | Names | Conf. % | Sequence |
| *Artemisia annua* | Actin-binding, cofilin/tropomyosin type | 99 | VGSPDESYEDFTNSLPSNECR |
|  | Actin-binding, cofilin/tropomyosin type | 99 | IEEQQVIVEK |
| *Artemisia sieversian* | Actin-binding, cofilin/tropomyosin type | 99 | IEEQQVIVEK |
|  | Actin-binding, cofilin/tropomyosin type | 99 | VGSPDESYEDFTNSLPSNECR |

**Figure 2.**
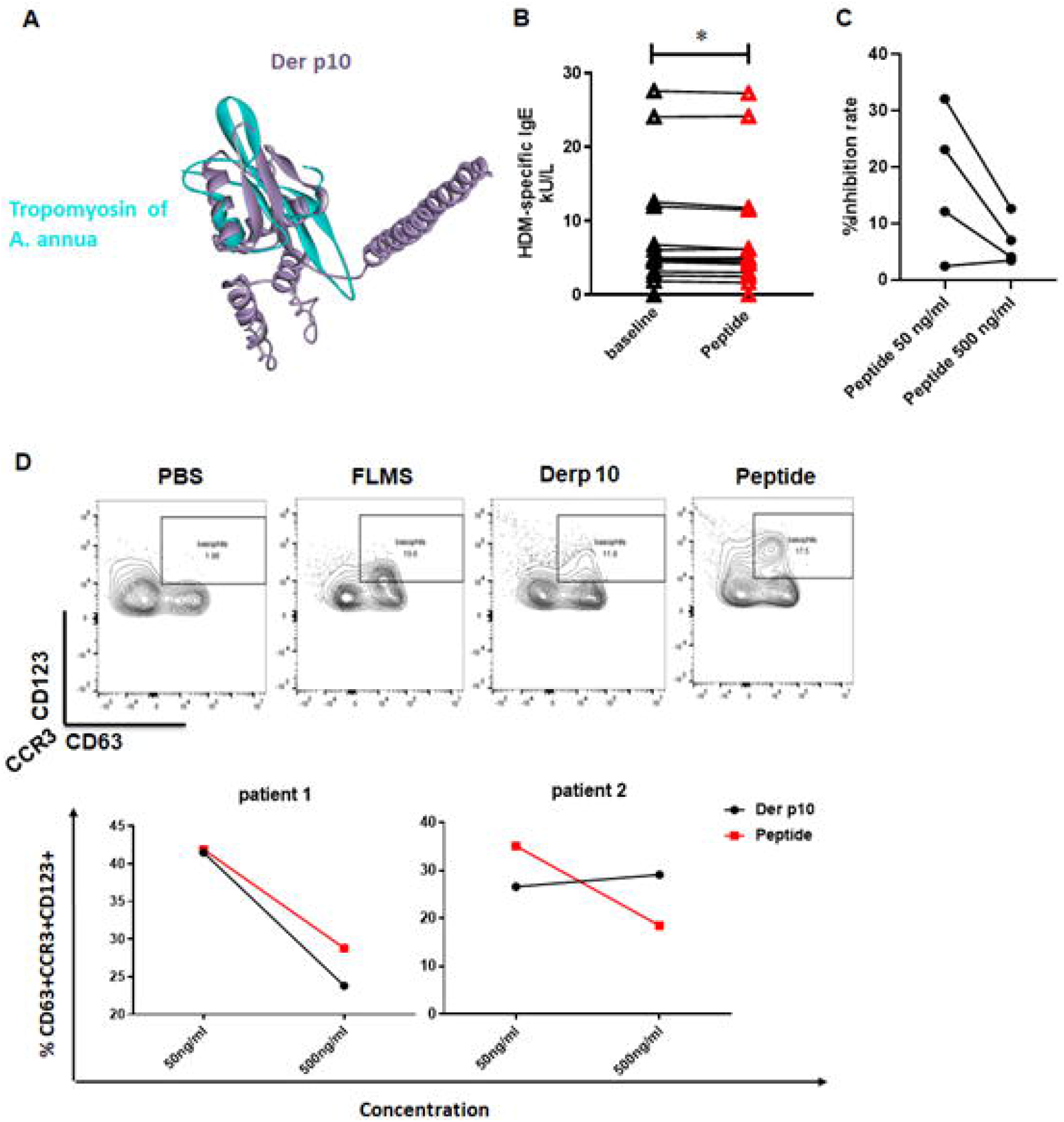
**A**: Structural comparison of tropomyosin of Artemisia annua (A. annua) and Dermatophagoides pteronyssinus 10 (Der p10). The target sequence of homology modelling tropomyosin of A. annua was established on the SWISS-MODEL server eEF1A2 (Swiss-Prot Accession No: Q05639). The amino acid sequences of each protein were then loaded into the SWISS-MODEL server to predict a 3D model. **B:** Concentrations of house dust mite (HDM)-specific IgE in the serum of HDM^+^ allergic rhinitis (AR) patients (n = 15) incubated in the absence or presence of synthesized tropomyosin peptides of mugwort; **C:** Tropomyosin of Artemisia annua (A. annua) peptide inhibits IgE-binding to Dermatophagoides pteronyssinus 10 (Der p10) measuring by ELISA. Tropomyosin of A. annua exhibits inhibitory effect in 4 out of 8 patients at the concentration of 50 ng/mL. **D:** The expression of basophils (CCR3^+^CD123^+^CD63^+^) with stimulations of Der p10 and tropomyosin of A. annua peptide stimulations. Totally, HDM specific IgE pre-incubated basophils from 2 out of 8 patients were activated by tropomyosin of A. annua.

Pre-incubation serum samples of monosensitized HDM^+^AR patients with mugwort tropomyosin peptide significantly decreased the concentrations of HDM specific-sIgE in the serum (**Figure 2B**). Furthermore, mugwort tropomyosin peptide (50 ng/ml) inhibited IgE binding to Der p10 ranging from 2.4% to 32.1% (**Figure 2C**). As shown in **Figure 2D**, the activation of basophils, which pre-sensitized by HDM specific IgE, occurred with exposure to Der p10 and synthetic tropomyosin mugwort peptide in 2 out of 8 non-allergic patients, suggesting the similar structure domains of tropomyosin in mugwort and HDM are functional.

### A. annua and synthetic tropomyosin of mugwort induced non-Th2 response in PBMCs of monosensitized HDM+AR patients

Compared to medium controls, HDM stimulated PBMCs isolated from monosensitized HDM^+^AR patients (n = 6) to produce high levels of IL-5 (0.59-61.56 pg/mL) and IL-17 (0.91-5.27 pg/mL), but not IFN-γ (**Figure 3A**). In contrast, HDM induced PBMCs isolated from healthy controls (n = 6) to release IL-17 (0.71-1.32 pg/mL) and IFN-γ (0.71-1.79 pg/mL), but not IL-5 (**Figure 3A**). Interestingly, A. annua stimulated PBMCs from monosensitized HDM^+^AR patients (n = 8) to produce IL-17 (1.36-2.24 pg/mL) and IFN-γ (3.7-4.79 pg/mL), but not IL-5 (**Figure 3B**); while such stimulation only induced the production of IFN-γ (1.79-47.45 pg/mL), but not IL-5 or IL-17 by PBMC of healthy controls (n = 5) (**Figure 3B**). Generally, the frequency of A. annua-induced release of IL-17 and IFN-γ was 25% and 12.5%, respectively, from PBMCs of monosensitized HDM^+^AR patients; and the frequency of HDM-induced release of IL-5 and IL-17 was 66.7% and 83.3%, respectively, from PBMCs of monosensitized HDM^+^AR patients **(Figure 3C)**, suggesting that HDM is more effective than A. annua in inducing inflammation in HDM^+^AR patients.

**Figure 3.**
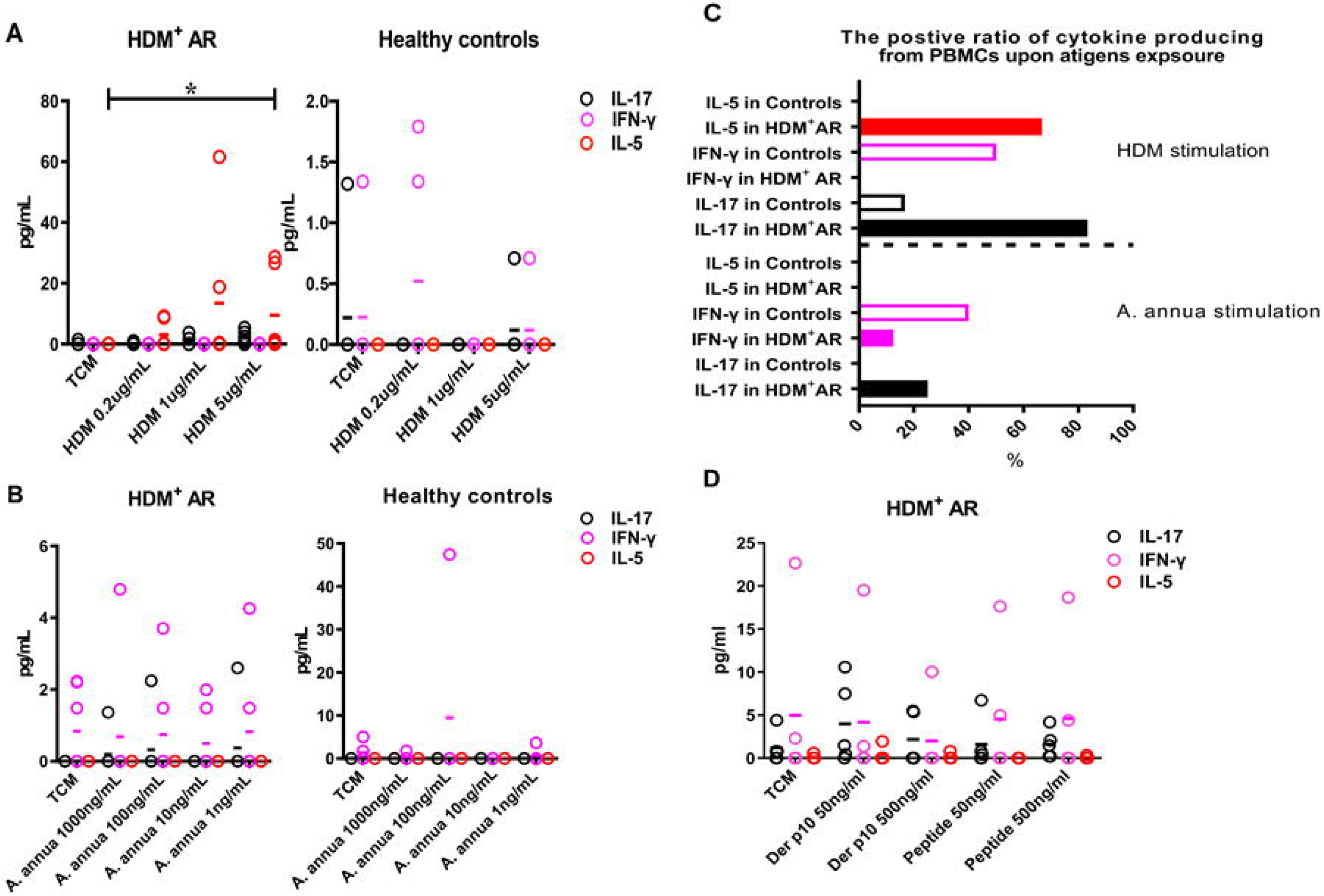
Effect of specific antigen on release of IL-17, IL-5 and IFN-γ in peripheral blood mononuclear cells (PBMCs) of HDM+ allergic rhinitis (AR) patients or healthy controls. **A:** Stimulation of PBMCs of monosensitized HDM^+^AR patients and control subject with house dust mite (HDM) (n = 6); **B:** Stimulation of PBMC of monosensitized HDM^+^AR patients (n = 8) and control subjects (n = 5) with Artemisia annua (A. annua); **C:** The positive ratio of cytokine produced by PBMCs of monosensitized HDM^+^ allergic rhinitis or controls upon HDM and A. annua stimulation. **D:** Concentrations of IL-17, IL-5 and IFN-γ released in PBMCs stimulated with synthesized tropomyosin peptides of A. annua and Der p10 (n = 4).

To confirm that tropomyosin may be responsible for A. annua extract-induced non-Th2 response in monosensitized HDM^+^AR patients, PBMCs from monosensitized HDM^+^AR patients were incubated with tropomyosin mugwort peptides and Der p10. Finally, mugwort peptide induced the synthesis and release of IFN-γ, with about 20% frequency and 60% of subjects released IL-17; whereas Der p10 induces the frequency of IL-5 and IL-17 were 20% and 60% respectively (**Figure 3D**).

## DISCUSSION

We demonstrated for the first time that cross-reactive protein tropomyosin in mugwort and HDM sharing similar structural domains is responsible for the cross-reactivity between HDM and mugwort. However, unlike HDM and Derp 10, A. annua and mugwort tropomyosin peptide induced Th1 and Th17 in PBMCs of HDM monosensitized AR patients, but not Th2 response.

Tropomyosin, a pan-allergen, belongs to a family of phylogenetically conserved proteins with multiple isoforms present in muscle and non-muscle cells of vertebrates and invertebrates^20^. It has been known that tropomyosin from HDM and cockroaches share high sequence homology with that of shellfish, which unsurprisingly results in cross-reactivity among HDM, cockroach and food allergens^21-23^. Mugwort is the most important outdoor seasonal allergen in Asia^4,24^. Our data have shown that a large number of polysensitized HDM^+^AR patients are sensitized to mugwort and the number of monosensitized HDM^+^AR patients was increased from July to August. Mite densities indeed vary with seasons and areas. Reportedly, three peaks for the domestic mites density in Beijing appear in September to October, January and May^25^. Thus, the increased number of monosensitized HDM^+^AR patients in July and August did be affected by mugwort. Although amino acid sequences of mugwort (A. annua and A. sieversian) tropomyosin proteins are different from that of HDM tropomyosin Der p10, 3D-modelling results show that the α-helices and β-sheets in mugwort tropomyosin are similar to Der p10. Considering that the sequence of the same protein varies in different species and cross-reactivity is thought to occur when a protein of similar sequence, structure or family binds to T and B cell receptors^26^, tropomyosin might be involved in mugwort-HDM cross-reactivity due to similar structure of tropomyosin in allergens.

The present study has indicated that stimulation of PBMCs of monosensitized HDM^+^AR patients with mugwort induced synthesis of IL-17 and IFN-γ, whereas stimulation with HDM induced synthesis of IL-5 and IL-17. Our group has previously demonstrated that single-nucleotide polymorphisms (SNPs) in IL-17A and IL-17F gene regions are potentially associated with the development of AR and comorbid asthma in Chinese subjects^27^. Similarly, a study in Caucasian subjects has also demonstrated that there is an association between serum IL-17 and the severity of clinical symptoms in AR patients^28^. As mentioned above, the role of Th17 in the pathogenesis of AR cannot be excluded. Thus, the induction of IL-17 by mugwort from PBMCs in monosensitized HDM^+^AR patients may be associated with clinical symptoms of patients. In this study, the finding for Der p10-induced synthesis of IL-5 in monosensitized HDM^+^AR patients was in accordance with the findings of stimulation by HDM. In the meanwhile, stimulation synthesized tropomyosin peptide of mugwort could induce IL-17 and IFN-γ by PBMCs of HDM^+^AR subjects, suggesting that tropomyosin is responsible for the cross-reactivity between HDM and mugwort, eliciting non Th2 response.

In conclusion, we have for the first time demonstrated that mugwort tropomyosin shares similar α-helices and β-sheets with Der p10, and therefore might play a role in eliciting a non-Th2 response in polysensitized HDM^+^AR patients in comparison to HDM. Since mugwort stimulation may be related to clinical symptoms of HDM sensitized patients, education and allergen avoid are required for HDM^+^AR patients in the autumn pollen season.

## Acknowledgements

This work was supported by grants from the Beijing Nova Program of Science and Technology (Z191100001119117); Yong top-notch talent (2018000021223ZK12); the Beijing Hospitals Authority Youth Programme (QML20180201); Beijing Scientific and Technological overall plan (Z171100000117002); the Beijing advanced innovation center for food nutrition and human health (Beijing Technology and Business University, 20181045); and the Program for the Changjiang Scholars and Innovative Research Team (IRT13082).

## REFERENCES

1. Akhouri S, House SA. Allergic Rhinitis. In: StatPearls. Treasure Island (FL): StatPearls Publishing Copyright © 2020, StatPearls Publishing LLC.; 2020.

2. Wang XD, Zheng M, Lou HF, et al. An increased prevalence of self-reported allergic rhinitis in major Chinese cities from 2005 to 2011. Allergy. 2016;71(8):1170–1180.

3. Wise SK, Lin SY, Toskala E, et al. International Consensus Statement on Allergy and Rhinology: Allergic Rhinitis. Int Forum Allergy Rhinol. 2018;8(2):108–352.

4. Lou H, Ma S, Zhao Y, et al. Sensitization patterns and minimum screening panels for aeroallergens in self-reported allergic rhinitis in China. Sci Rep. 2017;7(1):9286.

5. Matricardi PM, Dramburg S, Potapova E, Skevaki C, Renz H. Molecular diagnosis for allergen immunotherapy. J Allergy Clin Immunol. 2019;143(3):831–843.

6. Dogru M. Investigation of asthma comorbidity in children with different severities of allergic rhinitis. American journal of rhinology & allergy. 2016;30(3):186–189.

7. Tiotiu A, Plavec D, Novakova S, et al. Current opinions for the management of asthma associated with ear, nose and throat comorbidities. Eur Respir Rev. 2018;27(150).

8. Mahler V, Esch RE, Kleine-Tebbe J, et al. Understanding differences in allergen immunotherapy products and practices in North America and Europe. The Journal of allergy and clinical immunology. 2019;143(3):813–828.

9. Calderón MA, Cox L, Casale TB, Moingeon P, Demoly P. Multiple-allergen and single-allergen immunotherapy strategies in polysensitized patients: looking at the published evidence. The Journal of allergy and clinical immunology. 2012;129(4):929–934.

10. Li H, Chen S, Cheng L, et al. Chinese guideline on sublingual immunotherapy for allergic rhinitis and asthma. Journal of thoracic disease. 2019;11(12):4936–4950.

11. Bousquet J, Becker WM, Hejjaoui A, et al. Differences in clinical and immunologic reactivity of patients allergic to grass pollens and to multiple-pollen species. II. Efficacy of a double-blind, placebo-controlled, specific immunotherapy with standardized extracts. J Allergy Clin Immunol. 1991;88(1):43–53.

12. Kim KW, Kim EA, Kwon BC, et al. Comparison of allergic indices in monosensitized and polysensitized patients with childhood asthma. J Korean Med Sci. 2006;21(6):1012–1016.

13. Kim JY, Han DH, Won TB, et al. Immunologic modification in mono- and poly-sensitized patients after sublingual immunotherapy. Laryngoscope. 2019;129(5):E170–E177.

14. Cheng L, Chen J, Fu Q, et al. Chinese Society of Allergy Guidelines for Diagnosis and Treatment of Allergic Rhinitis. Allergy Asthma Immunol Res. 2018;10(4):300–353.

15. Sidenius KE, Hallas TE, Poulsen LK, Mosbech H. Allergen cross-reactivity between house-dust mites and other invertebrates. Allergy. 2001;56(8):723–733.

16. Brożek JL, Bousquet J, Agache I, et al. Allergic Rhinitis and its Impact on Asthma (ARIA) guidelines-2016 revision. The Journal of allergy and clinical immunology. 2017;140(4):950–958.

17. León IR, Schwämmle V, Jensen ON, Sprenger RR. Quantitative assessment of in-solution digestion efficiency identifies optimal protocols for unbiased protein analysis. Molecular & cellular proteomics: MCP. 2013;12(10):2992–3005.

18. Cantillo JF, Puerta L, Fernandez-Caldas E, et al. Tropomyosins in mosquito and house dust mite cross-react at the humoral and cellular level. Clin Exp Allergy. 2018;48(10):1354–1363.

19. Lou H, Huang Y, Ouyang Y, et al. Artemisia annua-sublingual immunotherapy for seasonal allergic rhinitis: A randomized controlled trial. Allergy. 2020.

20. Cantillo JF, Puerta L, Lafosse-Marin S, Subiza JL, Caraballo L, Fernandez-Caldas E. Allergens involved in the cross-reactivity of Aedes aegypti with other arthropods. Ann Allergy Asthma Immunol. 2017;118(6):710–718.

21. Valmonte GR, Cauyan GA, Ramos JD. IgE cross-reactivity between house dust mite allergens and Ascaris lumbricoides antigens. Asia Pacific allergy. 2012;2(1):35–44.

22. Santos AB, Chapman MD, Aalberse RC, et al. Cockroach allergens and asthma in Brazil: identification of tropomyosin as a major allergen with potential cross-reactivity with mite and shrimp allergens. J Allergy Clin Immunol. 1999;104(2 Pt 1):329–337.

23. Resch Y, Weghofer M, Seiberler S, et al. Molecular characterization of Der p 10: a diagnostic marker for broad sensitization in house dust mite allergy. Clin Exp Allergy. 2011;41(10):1468–1477.

24. Gao Z, Fu WY, Sun Y, et al. Artemisia pollen allergy in China: Component-resolved diagnosis reveals allergic asthma patients have significant multiple allergen sensitization. Allergy. 2019;74(2):284–293.

25. Sun JL, Shen L, Chen J, Yu JM, Yin J. Species diversity of house dust mites in Beijing, China. Journal of medical entomology. 2013;50(1):31–36.

26. M P, S H, R S, C D. Allergen cross-reactivity in allergic rhinitis and oral-allergy syndrome: a bioinformatic protein sequence analysis. International forum of allergy & rhinology. 2014;4(7):559–564.

27. Wang M, Zhang Y, Han D, Zhang L. Association between polymorphisms in cytokine genes IL-17A and IL-17F and development of allergic rhinitis and comorbid asthma in Chinese subjects. Human immunology. 2012;73(6):647–653.

28. Ciprandi G, De Amici M, Murdaca G, et al. Serum interleukin-17 levels are related to clinical severity in allergic rhinitis. Allergy. 2009;64(9):1375–1378.

